# Lactococcin B is inactivated by intrinsic proteinase PrtP digestion in *Lactococcus lactis* subsp. *lactis* BGMN1-501

**DOI:** 10.1101/309575

**Authors:** Goran Vukotic, Natalija Polovic, Nemanja Mirkovic, Branko Jovcic, Nemanja Stanisavljevic, Djordje Fira, Milan Kojic

**Author notes:** Goran Vukotic – Natalija Polovic - Nemanja Mirkovic - Branko Jovcic - Nemanja Stanisavljevic - Djordje Fira - Milan Kojic. Corresponding author: Milan Kojic, Laboratory for Molecular Microbiology, Institute of Molecular Genetics and Genetic Engineering, University of Belgrade, Vojvode Stepe 444/a, PO Box 23, 11010 Belgrade, Serbia.

## Abstract

In our previous study, we showed that PrtP is able to impair bacteriocin LcnB activity despite being produced by the same organism, and even if they were encoded by the same plasmid. However, exact cleavage site within LcnB bacteriocin, as well as the activity of the resulting peptides remained unknown. Here we further explored the interplay between these two proteins and defined, using mass spectrometry, that the hydrolysis occurs between the sixth and seventh amino acid on the N terminus of LcnB. Although it was suspected that the cleaved form of LcnB could retain some level of activity, both chemically synthesized and recombinant variant of truncated LcnB exhibited no antimicrobial activity. Wild type form of LcnB was recombinantly overexpressed using the same expression system, its antimicrobial activity was tested before and after the treatment with PrtP proteinase, and the degradation products were analyzed with reverse-phase high-pressure liquid chromatography. The results confirmed the inactivity of the truncated LcnB and additionally corroborated the PrtP cleavage site in LcnB bacteriocin.

**Importance:** Lactococcal enzyme PrtP, considered as a growth promoting factor, is involved in casein breakdown and enabling bacteria efficient growth in amino acids-poor, but protein-rich media. However, its interaction with bacteriocins was not known until recently. Bacteriocin LcnB can also be considered as growth promoting factor, since its known physiological role mirrors in preventing competing bacteria of reaching high growth densities. In this manuscript, we define the exact peptide bond inside the bacteriocin LcnB which is recognized by PrtP. This N-terminal removal of six amino acids completely inactivates the bacteriocin. The biological function of such action remains elusive. It is unexpected that in the same strain one enzyme inactivates a protein important for survival, unless it is some type of regulation.

## Introduction

Proteolytic system of lactic acid bacteria (LAB) occupied much attention since the 1980’s (1) onward, given its importance for efficient bacterial growth and the role it plays in industrial processes where these bacteria found immense applications. There is massive amount of data in literature, including some excellent reviews, regarding composition, physiology, biochemistry, diversity and genetics of proteolytic system in different LAB, especially in lactococci (2–4). Generally, lactococcal proteolytic system comprises: a) large cell-envelope proteinases (CEP), responsible for protein digestion and generation of oligopeptides; b) multiple oligopeptide transporters, ATP-binding cassette transporter proteins that mediate the uptake of proteinase-derived peptides; and c) intracellular peptidases, enzymes responsible for amino acid release from transported peptides. Historically, first discovered and most studied system is the one present in *Lactococcus lactis*. Strains of this species have a long tradition of application and importance in cheese, butter and other dairy production, making *L. lactis* one of the most exploited bacteria overall and its cell-envelope proteinase PrtP one of industrially most important enzymes. Although detailed functioning model of PrtP enzyme was made following vast experimental results, it turned out that there are ubiquitous differences in structure and function of PrtPs, even between closely related strains of *L. lactis*.

Bacteriocins are small ribosomally synthesized antimicrobial peptides, produced by some bacteria to kill competing ones. Likewise, they have been thoroughly studied, and a body of data is available on broad array of different bacteriocins (5–7). However, it seems that bacteriocin research still has not achieved its peak, since they are being growingly investigated as potential substitutes to antibiotics. Given their antimicrobial nature, and the lack of new and efficient antibiotics, they are recognized as one of the plausible means for combating the worldwide rise of antibiotic resistance, primarily for applications in food/feed preparation (8, 9). However, number of groups reported successful applications of bacteriocins in animal models of various infections (10–12).

In our previous study (13), we have shown that proteinase PrtP alters the bacteriocin LcnB activity of *L. lactis* subsp. *lactis* BGMN1-501. That discovery was instigated by the analysis of the *lcnB* gene promoter activity, during bacterial growth in media with rising concentrations of casitone. It was observed that activity of the *lcnB* gene promoter was decreasing, while at the same time antimicrobial activity of LcnB was increasing. It was concluded that some other mechanism is compensating for the silencing of the *lcnB* gene transcription. Since the transcription of the *prtP* gene is also medium dependent, but in opposing manner, we set to investigate the possible interplay between these two proteins. Namely, the possibility of LcnB being a substrate for PrtP was studied, both by making PrtP^-^ mutants and monitoring their bacteriocin activity, as well as by treating of LcnB by proteinase extract and tracking the same trait. It was shown that, although produced by the same strain, PrtP is capable of impairing bacteriocin LcnB, as its zones of growth inhibition were enlarged in PrtP^-^ mutant and diminished in the PrtP extracts treatment. Nevertheless, whether or not cleavage actually happens *in vivo* and at which position in LcnB remained to be unambiguously evidenced. The aim of the present study was to confirm proteolytic cleavage of LcnB by PrtP in BGMN1-501, establish the exact position where the cleavage occurs and determine how this process reflects on the antimicrobial activity of resulting molecules.

## 1. Materials and methods

### 1.1 Bacterial strains, growth mediums and conditions

The bacterial strains and plasmids used in this study are listed in Table 1. *Lactococcus lactis* subsp. *lactis* BGMN1-501 (14) was grown in M17 medium (Oxoid) supplemented with D-glucose (0.5% w/v) (GM17) at 30 °C. In addition, chemically defined medium (CDM) was made according to Miladinov et al. (15). *Escherichia coli* DH5α and ER2523, used for cloning and propagation of constructs, were grown in Luria-Bertani (LB) broth (16) aerobically at 37 °C, unless otherwise specified. Agar plates were made by adding 1.5% (w/v) agar (Torlak Belgrade, Serbia) to the liquid media. Ampicillin in concentration 100 μg/mL was used for selection and maintaining of transformants.

**Table 1.**
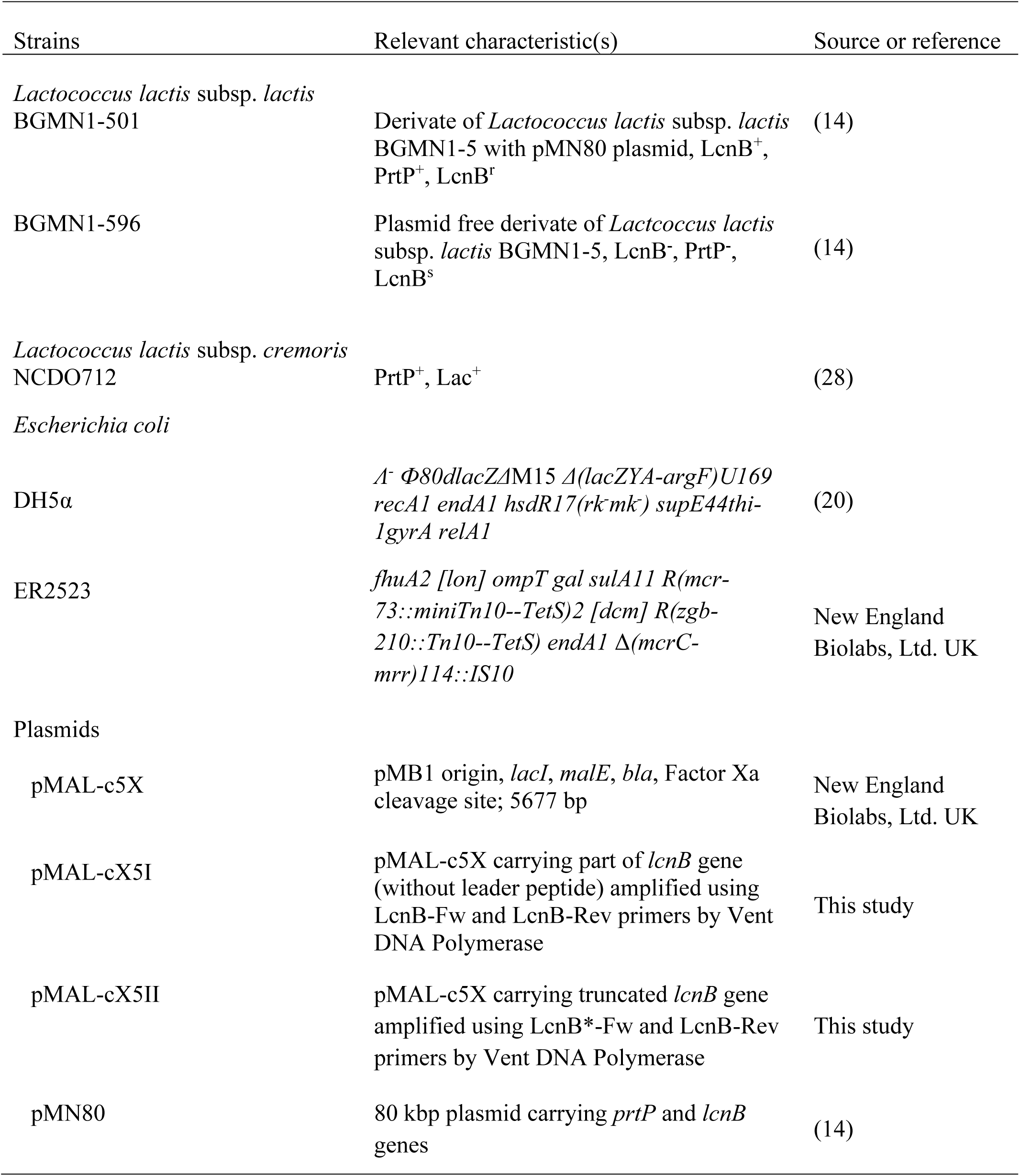
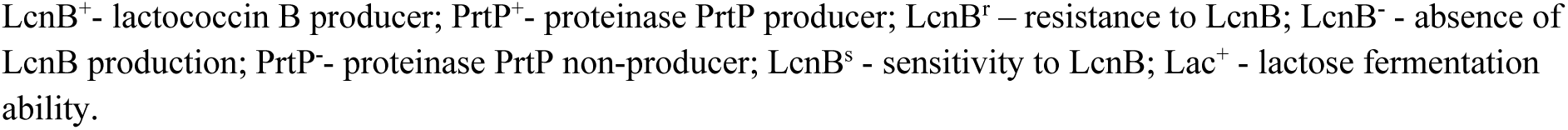
The bacterial strains and plasmids used in the study

### 1.2 Bacteriocin LcnB purification from *Lactococcus lactis* subsp. *lactis* BGMN1-501

For the purification and mass spectrometric analysis of the bacteriocin in *Lactococcus lactis* subsp. *lactis* BGMN1-501, the strain was grown at 30 °C for 16 h in 500 mL of chemically defined growth medium (CDM) with addition of 0.5% casitone (Difco Laboratories, Becton Dickinson and Company, Franklin Lakes, NJ, USA). The bacteriocin was purified following the protocol of Mirkovic et al. (17). Briefly, the cells were removed by centrifugation (4,500 × *g* for 30 min at 4 °C), while the supernatant was saturated up to 40% with ammonium sulfate, and precipitated with stirring for 2 h at 4 °C. The precipitate was collected by centrifugation (10,000 × *g* for 30 min at 4 °C) and then dissolved in 5 mL of Milli-Q water containing 0.1% trifluoroacetic acid. The bacteriocin was further purified using reverse-phase high-performance liquid chromatography (HPLC). Reverse-phase chromatography of the bacteriocin sample was performed using an Äkta Purifier 10 system (GE Healthcare, Uppsala, Sweden) with a Discovery BIO Wide Pore C_5_ column (10 cm by 4.6 mm; particle size, 5 μm; Supelco, Bellefonte, PA, USA). The protein was eluted using an acetonitrile gradient (0 to 90% with 0.1% trifluoroacetic acid for 10 column volumes). The chromatography was monitored by measuring absorbance at 215 nm. Twenty six obtained fractions were dried and dissolved in 10 μL of Milli-Q water, and tested for antibacterial activity.

### 1.3 Mass spectrometry analysis of antimicrobial active fraction

Mass spectrometry analysis of the active fraction was carried out using a mass spectrometer coupled with HPLC. The sample was injected onto a reverse-phase C18 column (RRHT column; 4.6 by 50 mm; particle size, 1.8 m) coupled with a Zorbax Eclipse XDB-C18 column installed in a 1200 series HPLC system (Agilent Technologies). The sample components were separated using an acetonitrile gradient (5 to 95% with 0.2% formic acid for 10 min and then 95% for 5 min). The mass spectrometer, a 6210 TOF liquid chromatography-mass spectrometry (LC-MS) system (G1969A; Agilent Technologies, Santa Clara, CA, USA), was run in positive ESI mode with a capillary voltage of 4.000 V, a fragmentor voltage of 200 V, and a mass range of m/z 100 to 3,200. Agilent MassHunter Workstation software and Analyst QS were used for data processing.

### 1.4 Bacteriocin activity

Bacteriocin activity of BGMN1-501, as well as of recombinantly produced bacteriocin was evaluated by an agar-well diffusion test (18).

The activity of HPLC-obtained fractions and of chemically synthesized bacteriocin was evaluated by spot-on-lawn assays. Soft GM17 agar (0.75%, w/v) containing sensitive *Lactococcus lactis* subsp. *lactis* BGMN1-596 was overlaid onto thin GM17 agar plates. After soft agar solidified, 10 μL of bacteriocin samples were spotted and plates were incubated at 30 °C overnight. Clear zones of growth inhibition in the spots were used as positive signals.

*In vitro* testing of effect of proteinase extract on bacteriocin activity was done as previously described (13), except that recombinant bacteriocins were used as substrates in the reactions.

### 1.5 DNA manipulations

Plasmid DNA from lactococci was isolated with the method described by O’Sullivan and Klaenhammer (19), while QIAprep Spin Miniprep Kit was used for plasmid DNA isolation from *E. coli,* according to the manufacturer’s recommendations (Qiagen, Hilden, Germany). Vent^®^ DNA Polymerase (New England Biolabs, Ltd. UK) was used to amplify DNA fragments by PCR, using GeneAmp PCR system 2700 thermal cycler (Applied Biosystems, Foster City, CA, USA). The primers used in PCR for amplification of the *lcnB* gene without leader peptide and its 18bp shorter truncated form, the *lcnB*\*, are listed in Table 2. Total DNA (1 ng) was mixed with 17.75 μL of bidistilled water, 2.5 μL of 10 × PCR buffer (Thermo Fisher Scientific), 1 μL dNTP mix (10 mM), 1.5 μL of MgCh (25 mM), 1 μL (10 pmol) of each primer and 0.25 μL of Vent^®^ DNA polymerase. PCR programs were consisted of initial denaturation (5 min at 96°C), 30 cycles of denaturation (30 s at 96°C), annealing (30 s at 40°C) and polymerization (1 min at 68°C), and an additional extension step of 5 min at 68°C. The PCR amplified DNA fragments were purified using QIAquick PCR Purification Kit as described by the manufacturer (Qiagen). T4 DNA ligase (Agilent Technologies, USA) was used for DNA ligation of PCR products into pMAL-c5X vector predigested with *Xmn*I, according to the manufacturer’s recommendation. Since Vent DNA Polymerase was used, blunt PCR amplicons were obtained, which was suited for ligation with pMAL-c5X vector, opened with *Xmn*I. Since blunt ligation offers two possible orientations, ligation mixes were transformed in DH5α competent cells, to screen for the appropriate fragment orientations. Standard heat-shock transformation was used for plasmid transfer into *E. coli* DH5α (20).

**Table 2.**
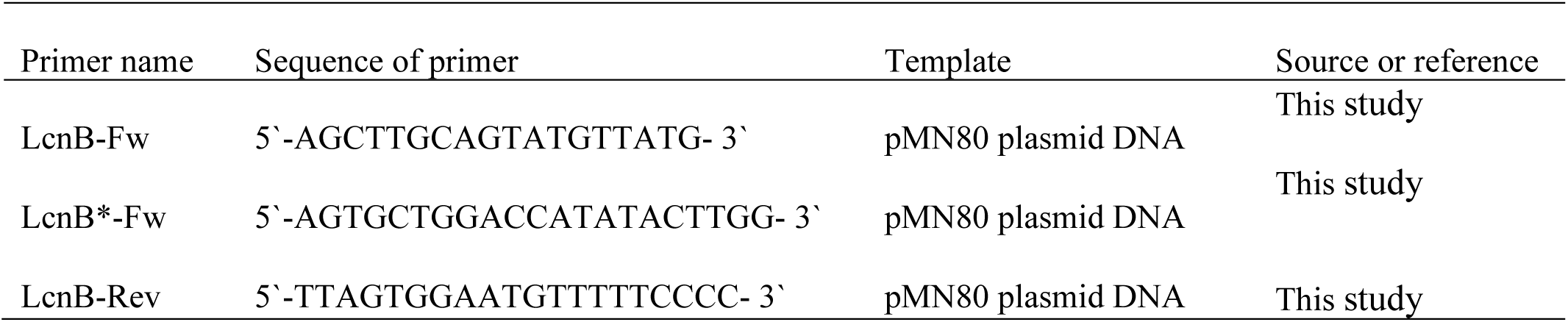
Sequence of specific primers used in this study.

Presence of cloned fragments in pMAL-c5X was determined using *Nco*I/*Sac*I digestion, while the orientation was determined by sequencing using Macrogen Sequencing Service (Macrogen, Netherlands). All digestions with restriction enzymes were conducted according to the supplier’s instructions (Thermo Fisher Scientific).

### 1.6 Chemical synthesis of bacteriocin

LcnB*, the truncated form of bacteriocin LcnB lacking first six amino acids (Table 3), was synthesized by Biomatik Corporation (Cambridge, Canada).

**Table 3.**
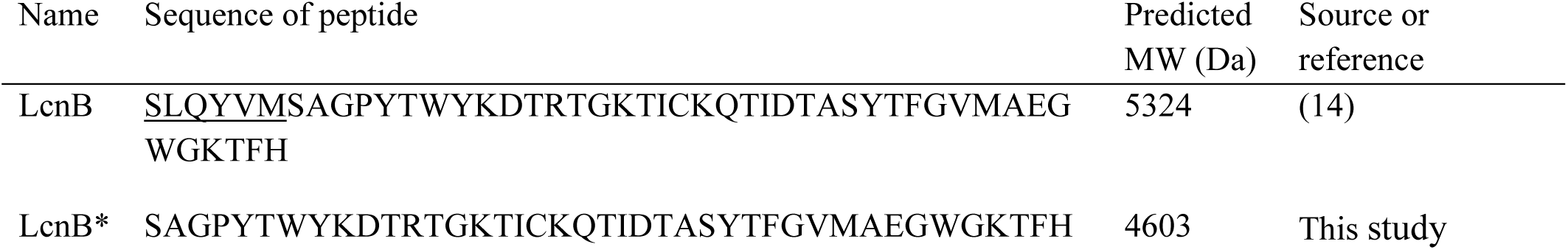
Amino acid sequences of bacteriocin forms used in this study. First six amino acids that are cleaved by PrtP proteinase are underlined in the wt LcnB bacteriocin.

### 1.7 Recombinant LcnB and LcnB* overexpression in *E. coli* ER2523 and purification

Recombinant proteins were overexpressed following the protocol of Miljkovic et al (21), with slight modifications. Plasmid constructs with appropriate fragment orientation, designated as pMAL-cX5I (carrying complete coding sequence for LcnB) and pMAL-cX5II (truncated for six amino acids at N terminus) were transformed into *E. coli* ER2523 competent cells (New England Biolabs). Transformants were selected on LA Petri dishes containing ampicillin (100 μg/mL) at 37 °C. Expression of recombinant proteins was carried out at 22°C by induction with 0.1 mM isopropyl β-D-1-thiogalactopyranoside (IPTG) for 2 h. Purification (cell lysis, affinity chromatography, cleavage of fusion protein with Xa protease) was performed according to manufacturer’s instructions (pMAL Protein Fusion & Purification System; New England Biolabs, Ltd. UK). Purified recombinant LcnB and LcnB* bacteriocins were stored at -20 °C in CM buffer (20 mM Tris-HCl pH 7.4, 200 mM NaCl, 1 mM EDTA, 1 mM DTT) containing 50% glycerol.

### 1.8 SDS-PAGE

Recombinant proteins were analyzed by SDS-PAGE, as previously described (13).

### 1.9 HPLC of recombinant proteins

Both recombinant proteins and the reaction mixtures of LcnB and PrtP were analyzed using the same reverse-phase chromatography as used for bacteriocin purification (section 2.2).

### 2.10 Bioinformatic analysis of LcnB

Bioinformatic analysis of LcnB and was carried out using JPred4, the latest version of JPred, one of the most accurate protein secondary structure prediction servers (22). This neural network-based server was used to predict propensity of each amino acid residue in given sequence to form certain type of secondary structure.

## 3. Results

### 3.1 Purification and mass spectrometric analysis of bacteriocin LcnB

BGMN1-501 was successfully cultivated in chemically defined medium and tested positively for bacteriocin activity (data not shown). The proteins present in the supernatant were precipitated by ammonium sulfate precipitation at 40% saturation, dialyzed and separated by RP-HPLC. Obtained fractions (twenty-six) were concentrated and tested for antimicrobial activity using spot-on-lawn method. The active fraction (fraction 24, Figure 1) was analyzed by ESI-TOF MS and the molecular mass of the present peptides was determined. Beside the peak of 5322 Da corresponding to predicted molecular mass of wild type LcnB (data not shown), additional peptide was detected (Figure 2). The mass analysis showed that the recorded molecular ion of this peptide was *m/z* 1534.8 ([M + 3H]^3+^), which indicated the molecular mass of 4601.4 Da. Based on LcnB amino acid sequence analysis, the detected peptide in the active fraction corresponded perfectly to theoretical LcnB molecular mass truncated of first six amino acids at its N terminus.

**Figure 1.**
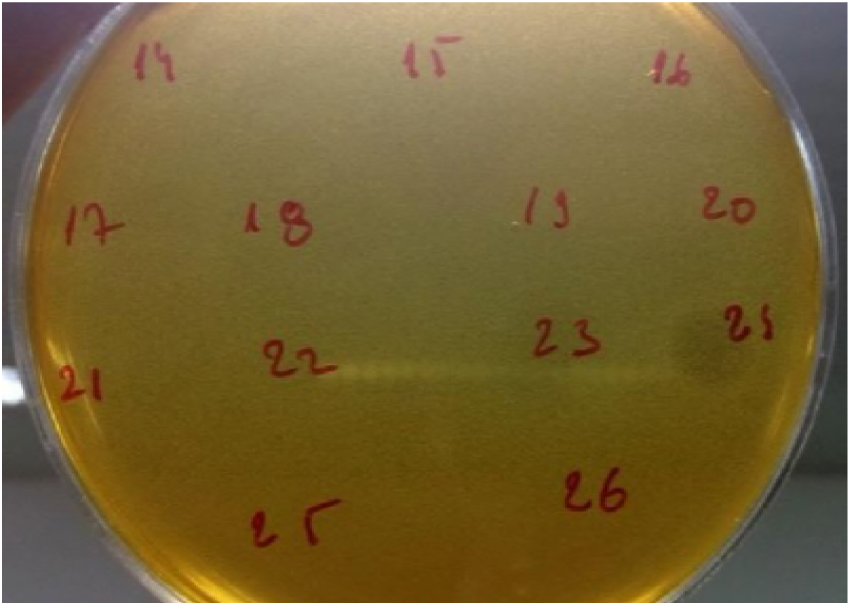
Testing of obtained fractions after HPLC purification of LcnB cleaved by PrtP proteinase for bacteriocin activity on *Lactococcus lactis* BGMN1-596.

**Figure 2.**
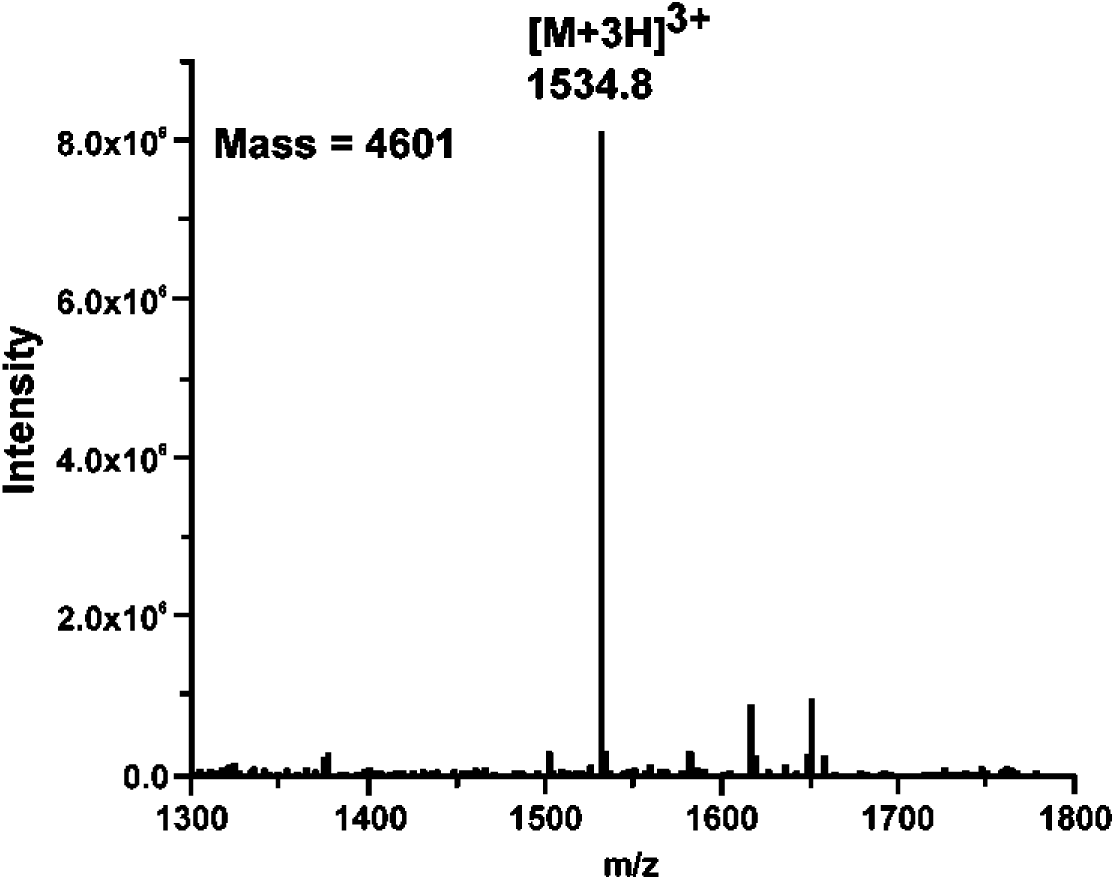
Mass spectrometry results of smaller peptide present in HPLC active fraction of LcnB cleaved by PrtP proteinase.

### 3.2 Truncated form of LcnB (LcnB*) is not active

When it was established that there are two proteins in the active fraction, we have set out experiment to determine and possibly compare the activity of both. For that reason, we ordered chemically synthesized truncated form of LcnB, shortened for six amino acids from N terminus, from Biomatik (Cambridge, Canada) and found it inactive on sensitive strain BGMN1-596, even in very high concentrations. However, since the manufacturer could not produce the wild-type peptide, we decided to produce both peptides by means of genetic engineering, in order to be sure in the obtained results and to have an adequate positive control. LcnB and LcnB* were produced as their mature peptide forms using pMAL protein fusion and expression system. Firstly, using plasmid pMN80 as template, both *lcnB* without leader peptide and its 18 bp shorter form were amplified using primer pairs LcnB-Fw/LcnB-Rev and LcnB*-Fw/LcnB-Rev, respectively. After determining the suitable clones, in which the fragments were cloned to be in-frame with maltose binding protein (MBP), two plasmids (named pMAL-cX5I, for matured LcnB and pMAL-cX5II for six amino acid shortened bacteriocin) were transformed into ER2523 strain for overexpression. Transformants were successfully obtained by selection on LA selective plates containing ampicillin (100 μg/mL) at 37 °C. Two transformants were selected and propagated at 22 °C for overexpression and purification of bacteriocin and its truncated form. The induction of expression under normal conditions (37 °C) resulted in insoluble proteins of interest. Similar results were also observed by other researchers (21, 23). Overview of the cloning and expression process is given in Figure 3.

**Figure 3.**
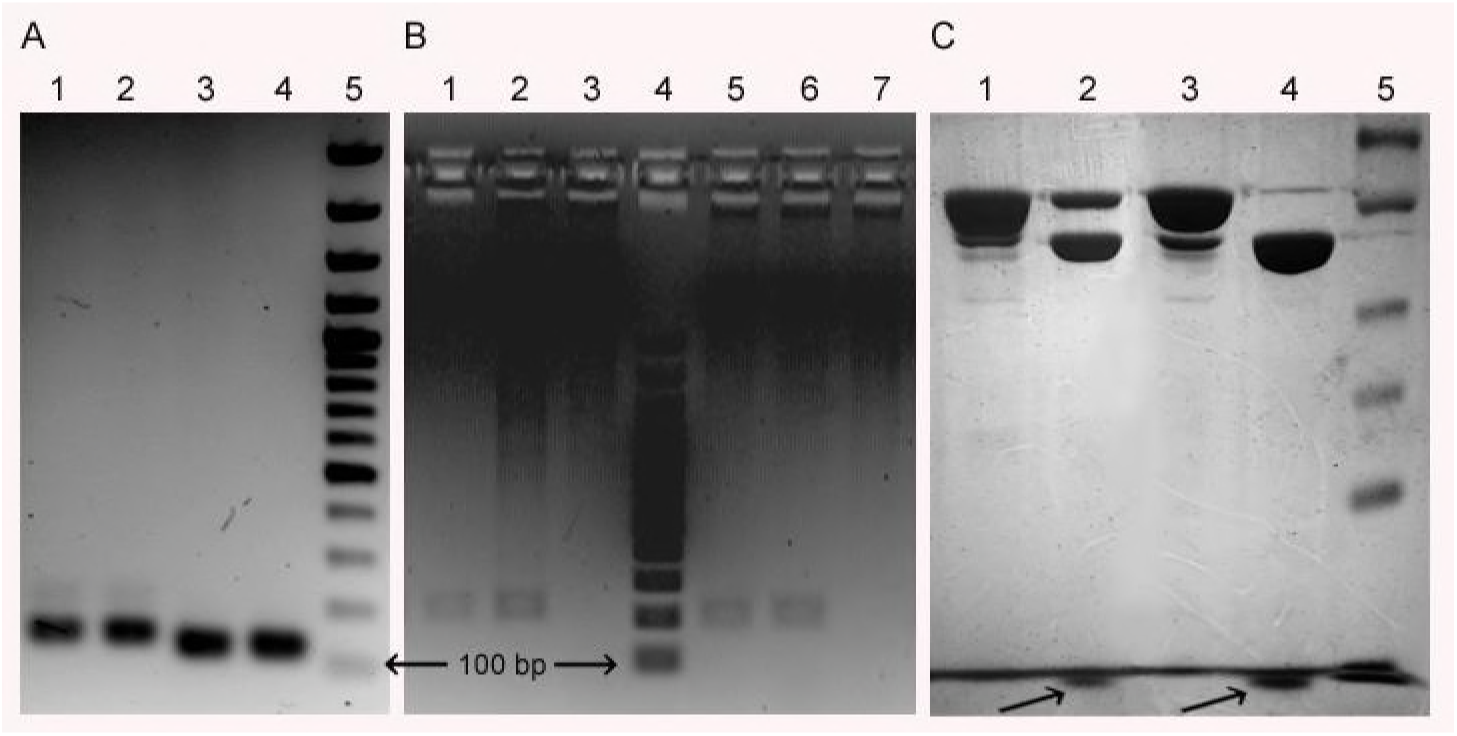
Production of LcnB and LcnB* by recombinant DNA technology. A) PCR amplification of *lcnB* (1, 2) and its truncated form (3, 4); 5, Generuler DNA Ladder Mix. B) *Nco*I/*Sac*I digestion of cloned PCR products in pMAL-c5X: 1-2, construct pMAL-c5XI (carrying *lcnB*); 5-6, construct pMAL-c5XII (carrying truncated form of *lcnB*); 3 and 7 empty vector; 4, Generuler DNA Ladder Mix. Note - the 90 bp increases in size compared to PCR amplicons is derived from vector. C) SDS-PAGE of overexpressed clones in *E. coli* ER2532: 1 and 3, fusions with MBP; 2 and 4, arrows pointing to proteins of interest (LcnB and LcnB*, respectively) released from MBP by Xa protease; 5 spectra multicolor broad range ladder.

When applied in spot-on-lawn bacteriocin activity assay, it was determined that LcnB possessed strong antimicrobial activity, corresponding to its concentration. However, LcnB* was proved to be inactive regardless of the concentration applied (Figure 4).

**Figure 4.**
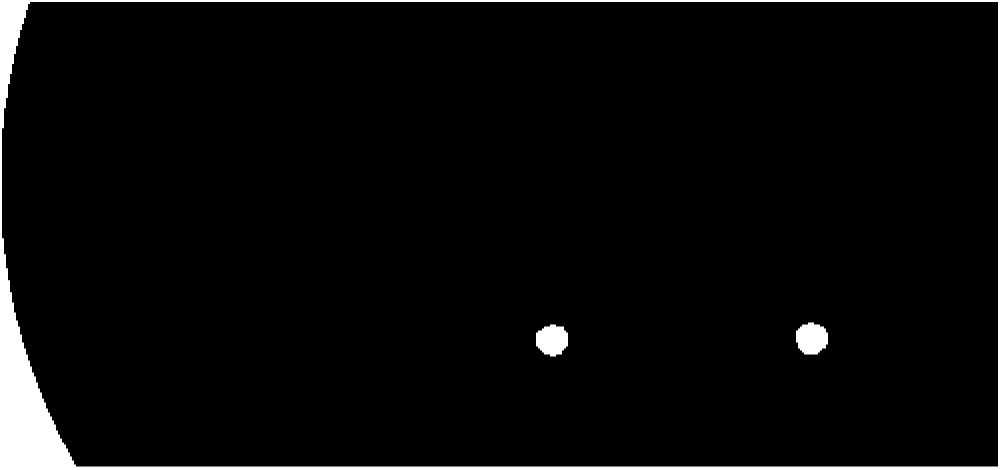
Antimicrobial activity of recombinant LcnB (A) and LcnB* (B). 10 μL of growing concentrations of each bacteriocin (the same concentrations of LcnB and LcnB* were applied) was spotted on lawn of sensitive cells *Lactococcus lactis* BGMN1-596. The most diluted sample was made by mixing 0.5 μL of bacteriocin preparation and 9.5 μL of sterile H_2_O. Labels correspond to the amount of bacteriocin (in μL) added to final 10 μL mix.

### 3.3 In vitro PrtP digestion of recombinant LcnB yields LcnB*

The definite confirmation of the previous results came with in vitro experiments where LcnB was subjected to PrtP hydrolysis as previously described (13). As expected, after 3 h of incubation PrtP reduced the bacteriocin activity of LcnB (Figure 5). Zones of growth inhibition, both in spot-on-lawn and agar-well test, diminished after the treatment with proteinase PrtP extract. However, some residual activity remains, which has been also demonstrated in our previous work on wild type bacteriocin, isolated from BGMN1-501. After 20 h of incubation with proteinase PrtP, no activity could be detected (data not shown).

**Figure 5.**
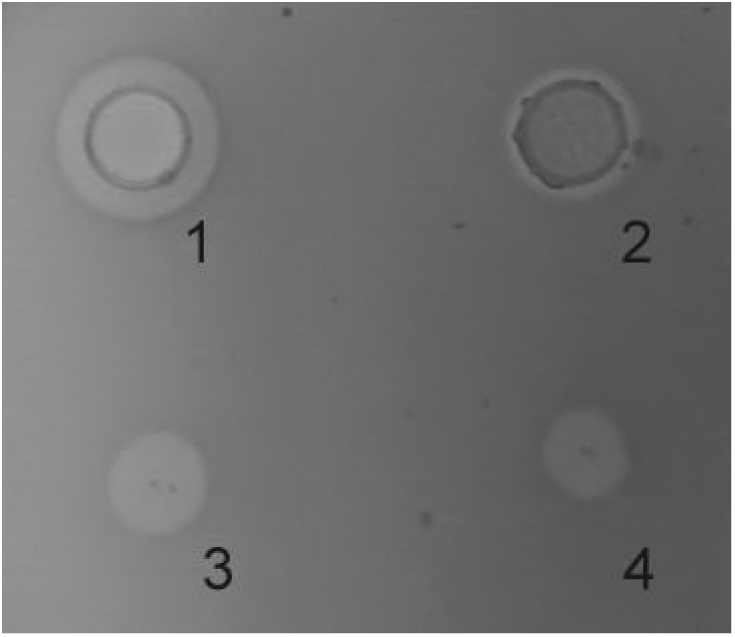
Antimicrobial activity of recombinant LcnB before and after 3 h incubation with PrtP proteinase in agar-well diffusion (1-2), and in spot-on-lawn (3-4) assays. 1 and 3, LcnB with NaPi buffer; 2 and 4, LcnB after 3 h of treatment with PrtP proteinase of *Lactococcus lactis* subsp. *cremoris* NCDO712 (reduced activity of LcnB was obtained). *Lactococcus lactis* BGMN1-596 was used as sensitive strain.

In order to investigate the pattern and kinetics of PrtP hydrolysis of LcnB, the digestion mix was separated using reverse-phase HPLC after 3 h and 20 h of digestion. In addition, recombinant LcnB as starting substrate and LcnB* as expected product were also loaded on the same column and separated under the same conditions, to establish the adequate controls (Figure 6). As can be seen on the figure, retention volume of wt LcnB is 11.9 mL in the given conditions (Figure 6, A), while for LcnB* this equals 12.2 mL (Figure 6, D). As for the 3 h hydrolysis (B), it can be seen that another peak is emerging in the close vicinity of the original one, perfectly corresponding to the peak generated by LcnB*. In addition, further degradation can be noted in the background. With the prolonged time of hydrolysis (Figure 6, C), it is clear that the amount of original substrate LcnB is drastically reduced, however, the same is happening to the LcnB* peptide – suggesting that apart from the specific digestion occurring at sixth amino acid in the protein, hydrolysis extends to other peptide bonds forming a background signal notable on the chromatogram.

**Figure 6.**
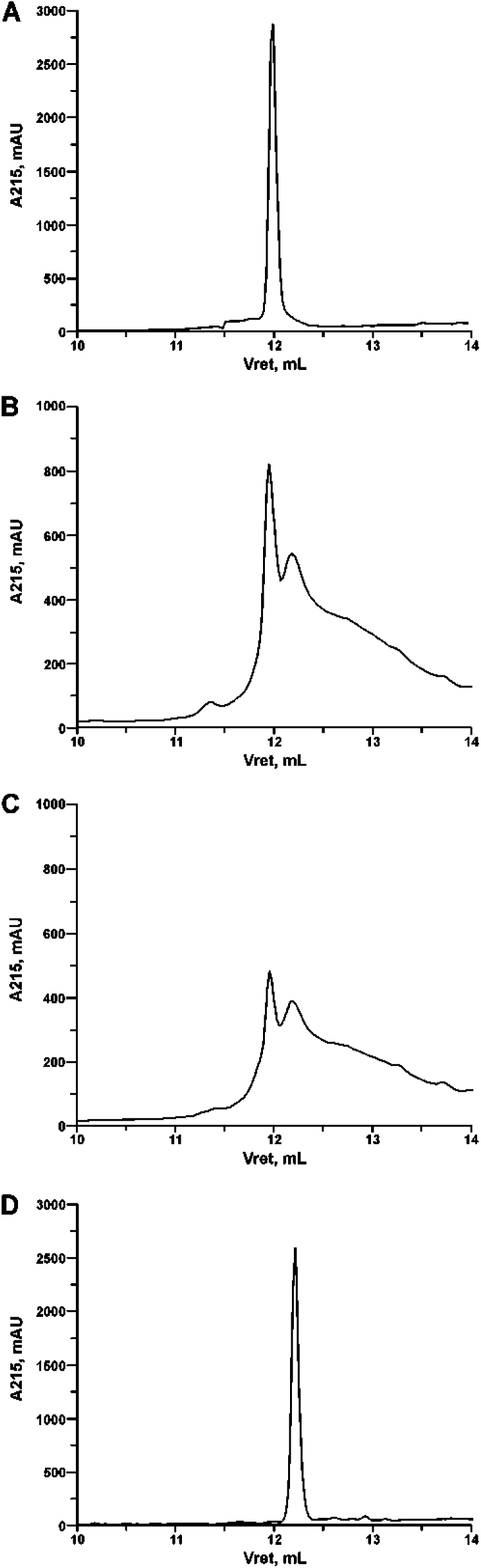
RF HPLC of recombinant LcnB (A), LcnB hydrolyzed with PrtP for 3 h (B), LcnB hydrolyzed with PrtP for 20 h (C) and recombinant LcnB* (D). Details are given in the text.

### 3.4 Bioinformatic analysis predicts loss of secondary structure in LcnB*

JPred4 analysis of amino acid sequence of wild type bacteriocin predicted three β-strands and one α-helix (Figure 7). The first β-strand comprises four residues from the first six amino acids (QYVM) of the protein, implying that this part of the peptide has an important function in secondary structure formation.

**Figure 7.**
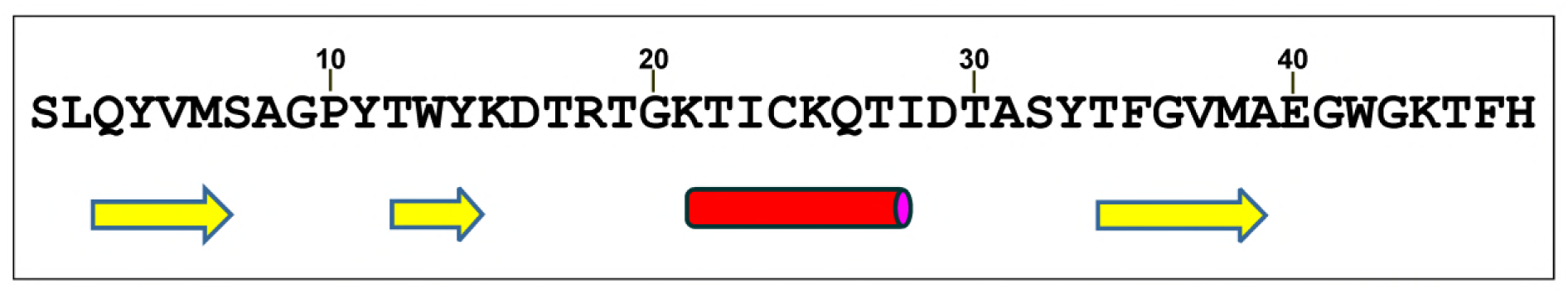
JPred4 prediction of secondary structure formation in LcnB. Yellow arrows: β-strands; Red cylinder: α-helix.

## 4. Discussion

In our previous paper, we described the medium dependent regulation of bacteriocin LcnB activity and gave the first insights into interaction between proteinase PrtP and LcnB bacteriocin. Through the generation and analyses of single-gene knockouts, it was shown that in BGMN1-501 bacteriocin activity is negatively regulated by PrtP, which is on the other hand negatively regulated by the peptide concentration in the growth medium. Finally, it was demonstrated that PrtP impaired LcnB activity in vitro, a result that fitted well and gave a plausible explanation for all recorded data. The same experiment indicated that proteinase might digest bacteriocin in a way that it remains partially active. However, regardless of how strong indications in vitro approach provided, an in vivo study was needed to confirm this unusual digestion, since it implies that BGM1-501, bacteriocin producer, decrease its own antimicrobial potential. This is why there was a need to determine whether hydrolysis actually occurs, at which peptide bond, as well as its severity to the function of bacteriocin. This was also interesting from the point of bacteriocin mode of action, given that this is still underexplored.

To approach this problem, we decided to purify and detect the present bacteriocin(s) from the overnight culture of a wild type bacterium BGMN1-501, by mass spectrometry. The rationale for this was the following: if the hydrolysis of LcnB does not occur, we would detect the bacteriocin as protein of adequate mass in the active fraction; if however, the hydrolysis of LcnB indeed occurs, we would either detect one protein of smaller mass or two proteins of different masses inside the active fraction, or two active fractions, each carrying an active protein.

To determine the form(s) of LcnB present in BGMN1-501 culture, the strain was grown in chemically defined minimal medium, supplemented with 0.5% casitone. This was done for two reasons: i) to grow bacteria in conditions favoring PrtP expression and hence putative bacteriocin digestion, and ii) to reduce the presence and number of molecules which could interfere with subsequent manipulations, i.e. chromatographic separation and detection of molecules of interest. The proteins present in the supernatant of such culture were separated with reverse-phase HPLC, generating 26 fractions. Active bacteriocin was clearly eluted in fraction 24, meaning it was one of the most hydrophobic proteins in the bacterial culture. Mass spectrometry revealed two proteins present in this fraction, one corresponding to theoretical bacteriocin LcnB, and, in addition, one of smaller mass. The protein sequence analysis revealed that the obtained 721 Da difference between the two detected peptides corresponded almost perfectly to theoretical value of first six amino acids in LcnB. This strongly indicated that the additional protein present in the active fraction is indeed the truncated form of LcnB protein.

This finding confirmed the hypothesis presuming that LcnB was actually hydrolyzed in the medium during bacterial growth. However, since the both peptides were detected in the active fraction, the activity of the truncated form, LcnB*, could not be ascertained. To define and compare the activity of both forms of bacteriocin, chemically synthesized proteins were ordered, but only truncated form was obtained. When tested for activity, this molecule seemed completely inactive. However, since without a positive control the definite conclusion could not be drawn, we decided to produce our own proteins of interest, using routine cloning work done in our lab. The pMAL expression system we used proved easy to handle, efficient and reliable, given that both proteins were expressed and purified without troubles. One of the positive features of this system is that it provides production of proteins exactly as desired by cloned DNA sequence–without the need for further processing.

Once purified, both proteins were tested for antimicrobial activity, both in spot-on-lawn and agar-well diffusion assays, however only LcnB was active, confirming the previous assumption that LcnB* does not possess bacteriocin activity. This was additionally corroborated with bioinformatic analysis of LcnB. Since detailed 3D structure of this bacteriocins is yet undetermined, the influence of given six amino acids removal on the final protein structure is hard to predict. However, secondary structure prediction suggested that these six amino acids are involved in β-sheet formation, implying the important function they have in protein structuring. Given that this β-sheet is predicted to be rather small, consisting of only 11 amino acids, it is highly probable that deletion of first β-strand, induced by PrtP, destabilize this structure completely, leading to denaturation of this β-sheet.

In order to test the susceptibility of recombinant LcnB to PrtP hydrolysis and with that to test all of our previous premises, an in vitro digestion was set up. PrtP extract obtained from NCDO712, a bacteriocin free strain, was mixed with LcnB and incubated for 3 h and 20 h in parallel. The reaction mixtures were tested by bacteriocin assays, and showed significant reduction in activity after 3 h (Figure 5), and no activity after 20 h (data not shown). These results were corroborated with HPLC analysis of pattern and kinetics of digestion, which gave final confirmation of our work. Namely, the 3 h chromatogram confirmed that PrtP digests LcnB, as original LcnB peak diminishes threefold, while another peak appears in its vicinity (Figure 6, A–B). Although severely reduced, there is still enough of starting LcnB left to produce visible growth inhibition (Figure 5, 2 and 4) of sensitive strain. The proteinase PrtP cleavage site is confirmed to be between 6^th^ and 7^th^ amino acid in the protein, as the retention volume of this additional peak coincides perfectly with the peak generated by recombinant LcnB* (Figure 6, D). It should also be noted that significant background signal appears in the 3 h chromatogram, which indicates additional hydrolysis. This is confirmed with the 20 h chromatogram (Figure 6, C) where both peaks are further diminished, although the LcnB* appears to be more stable (height of the peaks was almost the same after 20 h). This actually means that while LcnB* molecules are degraded with time, new ones are being generated through the activity of PrtP on LcnB, which keeps the pool of LcnB* in certain balance in first 20 h of digestion (Figure 6, B-C). Judging from the profile of background signal, it can be safely assumed that this additional hydrolysis is nonspecific. Therefore, it can be concluded that the preferred site for PrtP action is methionine-serine bond at the N terminus of LcnB, however, peptides liberated by this action will be hydrolyzed nonspecifically with time.

Taking all the results in consideration, we can conclude that our newest findings confirm hypotheses formed in our previous work, but not entirely. Proteinase PrtP indeed hydrolyzes bacteriocin LcnB, but by doing so, inactivates it. The reason for this eludes the common microbial logic. There seems to make no sense for bacteria to inactivate their own weapons for competition. However, risk of this exists only if PrtP is excessively expressed, which happens only when bacteria grow in nitrogen-depleted media, and are in other words starving. Nevertheless, it should be noted that conditions applied in this study mimicked such situations, given that bacteria were grown in peptide-poor chemically defined medium that induced PrtP expression, and yet, active bacteriocin was present in supernatant. We have already hypothesized that this “self-digestion” could have a physiological role in terms of obtaining of small peptides when they are extremely needed (13).

One should bear in mind that PrtP is being activated through the autoproteolytic removal of the N-terminal pro-region, during the maturation process. In addition, active enzymes are prone to additional auto-digestion, especially when no Ca^2+^ ions are present to stabilize the enzyme on the cell surface (24). Finally, PrtP is capable of hydrolyzing autolysin AcmA, a protein responsible for cell separation and autolysis during the stationary phase of growth (25), as well as some extracellular proteins of unknown function (24). Taken all together, it is possible that LcnB digestion by PrtP has some function, other than peptide generation, which still eludes our understanding. This is supported by the notion that LcnB degradation by PrtP is never so intense to completely deplete bacteria of their antimicrobial potential.

The results obtained in this study are important as they also proved that the first six amino acids are essential for bacteriocin function, and we presume that they constitute receptor-binding domain. This result are in agreement with previous theoretical considerations of Venema et al. (26) and experimental findings of Lasta et al. (27), who proposed methionine at position 6 of bacteriocin protein to be important for receptor binding in LcnB.

## Funding

This work was supported by the Ministry of Education and Science of the Republic of Serbia, Republic of Serbia (Grants No. 173019 and 173026).

## References

1. Law BA, Kolstad J. 1983. Proteolytic systems in lactic acid bacteria. Antonie Van Leeuwenhoek 49:225–45.

2. Pritchard GG, Coolbear T. 1993. The physiology and biochemistry of the proteolytic system in lactic acid bacteria. FEMS Microbiol Rev 12:179–206.

3. Law J, Haandrikman A. 1997. Proteolytic enzymes of lactic acid bacteria. Int Dairy J 7:1–11.

4. Savijoki K, Ingmer H, Varmanen P. 2006. Proteolytic systems of lactic acid bacteria. Appl Microbiol Biotechnol 71:394–406.

5. Diep DB, Nes IF. 2002. Ribosomally synthesized antibacterial peptides in Gram positive bacteria. Curr Drug Targets 3:107–22.

6. Cotter PD, Ross RP, Hill C. 2013. Bacteriocins - a viable alternative to antibiotics? Nat Rev Microbiol 11:95–105.

7. Gabrielsen C, Brede DA, Nes IF, Diep DB. 2014. Circular bacteriocins: biosynthesis and mode of action. Appl Environ Microbiol 80:6854–62.

8. Cotter PD, Hill C, Ross RP. 2005. Bacteriocins: developing innate immunity for food. Nat Rev Microbiol 3:777–88.

9. Mills S, Ross RP, Hill C. 2017. Bacteriocins and bacteriophage; a narrow-minded approach to food and gut microbiology. FEMS Microbiol Rev 41:S129–S153.

10. Mota-Meira M, Morency H, Lavoie MC. 2005. In vivo activity of mutacin B-Ny266. J Antimicrob Chemother 56:869–71.

11. Dabour N, Zihler A, Kheadr E, Lacroix C, Fliss I. 2009. In vivo study on the effectiveness of pediocin PA-1 and Pediococcus acidilactici UL5 at inhibiting Listeria monocytogenes. Int J Food Microbiol 133:225–33.

12. Heunis TDJ, Smith C, Dicks LMT. 2013. Evaluation of a nisin-eluting nanofiber scaffold to treat Staphylococcus aureus-induced skin infections in mice. Antimicrob Agents Chemother 57:3928–35.

13. Vukotic G, Mirkovic N, Jovcic B, Miljkovic M, Strahinic I, Fira D, Radulovic Z, Kojic M. 2015. Proteinase PrtP impairs lactococcin LcnB activity in Lactococcus lactis BGMN1-501: new insights into bacteriocin regulation. Front Microbiol 6:92.

14. Kojic M, Strahinic I, Fira D, Jovcic B, Topisirovic L. 2006. Plasmid content and bacteriocin production by five strains of Lactococcus lactis isolated from semi-hard homemade cheese. Can J Microbiol 52:1110–20.

15. Miladinov N, Kuipers OP, Topisirovic L. 2001. Casitone-mediated expression of the prtP and prtM genes in Lactococcus lactis subsp. lactis BGIS29. Arch Microbiol 177:54–61.

16. Miller JH. 1972. Experiments in molecular genetics.

17. Mirkovic N, Polovic N, Vukotic G, Jovcic B, Miljkovic M, Radulovic Z, Diep DB, Kojic M. 2016. Lactococcus lactis LMG2081 Produces Two Bacteriocins, a Nonlantibiotic and a Novel Lantibiotic. Appl Environ Microbiol 82:2555–62.

18. Lozo J, Vukasinovic M, Strahinic I, Topisirovic L. 2004. Characterization and antimicrobial activity of bacteriocin 217 produced by natural isolate Lactobacillus paracasei subsp. paracasei BGBUK2-16. J Food Prot 67:2727–34.

19. O’sullivan DJ, Klaenhammer TR. 1993. Rapid Mini-Prep Isolation of High-Quality Plasmid DNA from Lactococcus and Lactobacillus spp. Appl Environ Microbiol 59:2730–3.

20. Hanahan D. 1983. Studies on transformation of Escherichia coli with plasmids. J Mol Biol 166:557–80.

21. Miljkovic M, Malesevic M, Filipic B, Vukotic G, Kojic M. 2018. LraI from Lactococcus raffinolactis BGTRK10-1, an Isoschizomer of EcoRI, Exhibits Ion Concentration-Dependent Specific Star Activity. Biomed Res Int 2018:1–10.

22. Drozdetskiy A, Cole C, Procter J, Barton GJ. 2015. JPred4: a protein secondary structure prediction server. Nucleic Acids Res 43:W389–94.

23. Chung D-H, Huddleston JR, Farkas J, Westpheling J. 2011. Identification and characterization of CbeI, a novel thermostable restriction enzyme from Caldicellulosiruptor bescii DSM 6725 and a member of a new subfamily of HaeIII-like enzymes. J Ind Microbiol Biotechnol 38:1867–77.

24. Haandrikman AJ, Meesters R, Laan H, Konings WN, Kok J, Venema G. 1991. Processing of the lactococcal extracellular serine proteinase. Appl Environ Microbiol 57:1899–904.

25. Buist G, Venema G, Kok J. 1998. Autolysis of Lactococcus lactis is influenced by proteolysis. J Bacteriol 180:5947–53.

26. Venema K, Dost MH, Venema G, Kok J. 1996. Mutational analysis and chemical modification of Cys24 of lactococcin B, a bacteriocin produced by Lactococcus lactis. Microbiology 142 (Pt 10):2825–30.

27. Lasta S, Fajloun Z, Mansuelle P, Sabatier JM, Boudabous A, Sampieri F. 2008. [Chemical synthesis of lactococcin B and functional evaluation of the N-terminal domain using a truncated synthetic analogue]. Arch Inst Pasteur Tunis 85:9–19.

28. Gasson MJ. 1983. Plasmid complements of Streptococcus lactis NCDO 712 and other lactic streptococci after protoplast-induced curing. J Bacteriol 154:1–9.

